# *NLR1-V*, a CC-NBS-LRR encoding gene, is a potential candidate gene of the wheat powdery mildew resistance gene *Pm21*

**DOI:** 10.1101/114058

**Authors:** Liping Xing, Ping Hu, Jiaqian Liu, Chaofan Cui, Hui Wang, Zhaocan Di, Shuang Zhou, Jiefei Xu, Li Gao, Zhenpu Huang, Aizhong Cao

## Abstract

Wheat powdery mildew caused by *Blumeria graminisb* f. sp. *tritici* is one of the most destructive diseases all over the world. *Pm21,* transferred from the wild *Haynaldia villosa* to wheat, confers broad spectrum resistance throughout the whole stage, and this gene has been widely used in wheat production for more than 20 years. Cloning the candidate gene of *Pm21* is the prerequisite for elucidating the resistance mechanism, and is a valuable attempt to clone the target genes from the evolutionarily distant wild species. In this study, an innovative approach, which combined cytogenetic stocks development, mutagenesis, RenSeq and PacBio, was tried successfully to clone an NBS-LRR type gene *NLR1-V* from the *Pm21* locus. Firstly, a powdery mildew resistant cryptic alien introgression line HP33 involved very small 6VS segment was developed, and 6 independent susceptible mutants of T6VS · 6AL was identified. Then, the transcriptome of *H. villosa* was obtained by NGS and the full-length NBS-LRR gene database was constructed by RenSeq-PacBio. In the following study, two expressed *NLR* genes were located to the *Pm21* locus using the HP33 as the mapping material, and only *NLR1-V* showed polymorphism between the wild T6VS · 6AL and its six mutants. The functional analysis indicated that silencing of *NLR1-V* could compromise the resistance of T6VS · 6AL completely, and could also decrease the resistance of T6VS · 6DL dramatically. Moreover, *NLR1-V* could recover the resistance of the susceptible mutant and increase the resistance in the susceptible wheat. The study implied that *NLR1-V,* a CC-NBS-LRR encoding gene, is a potential candidate gene of the powdery mildew resistance gene *Pm21.*

## Introduction

Plant faces numerous pathogens during their whole growth stage, and has evolved sophisticated mechanisms to escape the damages induced by different diseases, including the passive resistance pattern and the active resistance pattern. The passive resistance was usually mediated by the physical barriers on the surface of the plant, and this type of resistance can help plants protect themselves from most of the pathogens. The active resistance pattern involved two types of receptors, one type of receptors named as pattern-recognition receptors (PRR) which recognize the pathogen-associated molecular patterns (PAMP), and this type of receptors induced the resistance as PTI (PAMP triggered resistance). The other type of receptors, usually referred to as the resistance gene (R gene), could recognize the specific effectors produced by pathogens, and this type of receptors induced the resistance as ETI (effector triggered resistance). The resistance of PTI is usually possessed by all genotypes of one plant species, however, the resistance of ETI is usually possessed by the specific genotype of one plant species (Maekawa et al., 2011).

More than 100 R genes have been cloned from various species, and most of them contained two conserved domains, including nucleotide-binding sites and leucine-rich repeat (NBS-LRR, NLR) (Yang et al., 2013). According to the conserved domain in the N terminal, the cloned R genes were classified into two types, one type contains the conserved domain resembling the Drosophila Toll and mammalian IL-1 receptors (TIR) and named as TIR subclass, and the other named as non-TIR class (Meyers et al. 1999). In the non-TIR class, most NLR genes contain the coiled-coil (CC) domain, and a few NLR genes contain the leucine zipper motif or RPW8 domain (Gu et al., 2015). The genomic data of a number of species have been sequenced, and it was found that dicot species contained both the TIR class and non-TIR class, while the monocot species contained only the non-TIR class. For example, in the Arabidopsis 67% NLR genes were TIR class and 33% NLR genes were non-TIR class among the 159 NLR genes (Guo et al. 2011), however, in the rice all the 535 NLR genes were belonging to the CC-NLRs (Zhou et al. 2004).

Up to date, most cloned R genes in the cereals were characterized as CC-NLRs, such as *Mla1*(Zhou et al.2001), *Mla6* (Halterman et al., 2001), *Mla10* (Halterman and Wise 2004), *Mla12* (Shen et al., 2003), *Mla13* (Halterman et al., 2003) and *Rpg5* (Arora et al., 2013) in barley, *Pita* (Wang et al. 1999), *Pib* (Bryan et al. 2000), *Pi36* (Liu et al. 2007b), *Pi64* (Ma et al., 2015) and *PigmR* (Deng et al., 2016) in rice, *Rp1*(Collins et al. 1999) in maize. As in wheat, except the *Lr34* (Krattinger et al., 2009), *Yr36* (Fu et al., 2009) and *Lr67* (Moore et al., 2015), al the other cloned R genes were all belonging to the CC-NLRs, including the *Lr1* (Cloutier et al., 2007), *Lr10* (Feuillet et al., 2003) and *Lr21* (Huang et al., 2009) conferring resistance to leaf rust, *Yr10* (Liu et al., 2014) conferring resistance to stripe rust, *Sr22* (Steuernagel et al., 2016), *Sr33* (Periyannan et al., 2013), *Sr35* (Saintenac et al., 2013), *Sr45* (Steuernagel et al., 2016) and *Sr50* (Mago et al., 2015) conferring resistance to stem rust, and *Pm2* (Sanchez-Martin et al., 2016), *Pm3* (Yahiaoui et al., 2009) and *Pm8* (Hurni et al., 2013) conferring resistance to powdery mildew.

Wheat powdery mildew, caused by *Blumeria graminisb* f. sp. *tritici* (*Bgt*), is one of the most destructive diseases of wheat globally especially in cool and humid areas, and it has been reported to induce severe grain yield losses, ranging from 13 to 34% (Griffey et al. 1993;). The most environmental friendly and economical way to reduce the losses caused by powdery mildew disease in wheat is to develop varieties with high resistance (Kuraparthy et al. 2007). To date, more than 73 genes/alleles conferring resistance to powdery mildew have been identified at 50 gene loci in wheat and its relatives (Petersen et al. 2015), however, most of which confer isolate specific resistance. *Bgt* is a kind of pathogen with high evolutionary speed leading to the rapid isolates variation, and new virulent isolates usually emerged following the widely deployment of resistance genes especially those isolate-specific resistance genes. Durable and broad-spectrum resistance genes is usually sustainable which cannot be easily conquered by the new virulent isolates, so it is valuable to explore such kind of genes to counter the rapid evolution of *Bgt* populations and promote their use in the crop production.

*Haynaldia villosa* (2n=2x =14, VV), a diploid wheat wild relative species, contains powdery mildew resistance gene *Pm21*, which was a durable and broad-spectrum resistance gene conferring high resistance to more than 120 isolates in both China and Europe (Huang et al., 1997). *Pm21* was introduced from *H. villosain* to common wheat by developing the whole-arm translocation T6VS/6AL (Chen et al., 1995), which has been used as the parent of more than 30 varieties cultivated in about four million hectares in different wheat-growing areas in China (Bie et al. 2015), and *Pm21* has become one of the most highly effective genes introgressed into wheat from its wild relatives up to date. *Pm21* was firstly located on the 6VS arm (Chen et al. 1995), and subsequently located in the region of FL 0.45–FL 0.58 (Cao et al. 2011) by using a resistant deletion stock del.6VS-1 (FL0.58) and a susceptible deletion stock del.6VS-2 (FL0.45) (Cao et al., 2011). Chen (2013) then located *Pm21* at the region of CINAU273-CINAU276 using two translocation lines T6VS-2 and T6VS-1. However, these lines were identified by GISH, and the segment to locate the *Pm21* gene was still too large for the gene fine mapping. Different approaches have been tried to clone the *Pm21* gene, and *Stpk-V, DvMPK1, DvMLPK, DvUPK* and *DvPSYR1* were found to be related to the resistance mediated by 6VS (Cao et al., 2011; He et al., 2016). However, silent of these five genes can not compromised the resistance of T6VS · 6AL completely. Based on the reports that most of the cloned R genes in the cereal crops were belonging to the NLR class, it was hypothesized that if there is some NLR genes in the *Pm21* locus play even more important roles in the powdery mildew resistance.

In recent two years, new strategy combing the whole genome NLR gene exploration and mutant screening was used to clone the R genes in wheat, and several R genes were cloned rapidly and successfully (Periyannan et al., 2013; Mago et al., 2015; Steuernagel et al., 2016). In this study, a *NLR1-V* gene was cloned in the *Pm21* locus by NLR gene precisely location and functional analysis. Firstly, a new cryptic alien introgression line resistant to powdery mildew involved the very small 6VS chromosome segments was screened from a large irradiation population, and several individual susceptible mutants were identified from a big mutated population of T6VS · 6AL. At the same time, the NLR database of the whole genome of *H. villosa* was constructed by using RenSeq-Pacbio. Then, a *NLR1-V* gene, which was screened in the NLR database and located in the 6VS small segment in the resistant cryptic alien introgression line, was found to be responsible for the complete resistance to powdery mildew in the T6VS · 6AL. Mutated of *NLR1-V* gene could make T6VS · 6AL lose the powdery mildew resistance completely, and over-expression of it can recover the resistance in the susceptible mutant. So, it is proposed that this *NLR* gene *NLR1-V* could be an even more important gene than the ever explored resistance related genes to regulate the powdery mildew resistance in the *Pm21* locus, and it is the most likely candidate gene of *Pm21.*

## Materials and Methods

### Powdery mildew resistant wheat-*H*, *villosa* translocation line T6VS · 6AL and T6VS · 6DL

The wheat-*H*, *villosa* translocation line T6VS · 6AL was developed by Cytogenetic Institute of Nanjing Agricultural University (CINAU), in which the 6AS chromosome arm of wheat was substituted by the 6VS chromosome arm of *H.villosa* (Chen et al., 1995). T6VS · 6AL showed high and broad spectrum resistance to powdery mildew for the *Pm21* gene located in the 6VS chromosome arms. T6VS · 6DL was a wheat-*H*. *villosa* translocation line developed by Chinese Academy of Agricultural Sciences (CAAS), in which the 6DS of wheat was substituted by 6VS of *H.villosa* and the powdery mildew resistance gene *PmV* was located on the 6VS. The *H.villosa* donor of T6VS · 6AL was introduced from the United Kingdom but *H.villosa* donor of T6VS · 6DL was introduced from the Soviet Union, and diversity was detected between the two 6VS chromosome arms by molecular marker analysis (Bie et al., 2015). In this study, T6VS · 6AL was used as the wild line to compare with its mutants as aspect of the sequence of the candidate NLR genes, and both T6VS · 6AL and T6VS · 6DL were used for the BSMV inoculation for gene silencing to evaluate the effects of the NLR genes. *T. durum-H. villosa* amphiploid (AABBVV) and Chinese Spring (AABBDD) were developed or preserved by CINAU.

### A powdery mildew resistant cryptic alien introgression line HP33

To create the resistant introgression lines involved small 6VS segments, spikes of the T6VS · 6AL translocation line were irradiated 2–3 days before flowering with ^60^C_O_ γ-rays at a 160 Rad/M dosage rate, then the irradiated spikes were pollinated with common wheat after 2–3 days to produce the M1 seeds. The propagated plants were then identified by molecular markers and resistance evaluation. A new cryptic alien introgression, named HP33, was screen from the large irritation population which showed powdery mildew throughout the whole growth stage.

### Powdery mildew susceptible mutants of T6VS · 6AL induced by EMS

As described by Periyannan et al (2013), about 20,000 seeds of the T6VS.6AL translocation line carrying *Pm21* were treated with 0.7% EMS and about 3,500 independent plants were obtained at the M1 generation. Then they were advanced to the M2 generation and screened for response to *Bgt* race E31 (incompatible to *Pm21*). Six individual susceptible mutants (SM-1 to SM-6) derived from different M2 lines were screened and then were reconfirmed in the M3 generation. SM-1 and its wild type T6VS · 6AL were inoculated with 24 different *Bgt* isolates in CAAS, and the T6VS · 6AL showed the highest resistance level but the SM-1 showed the highest susceptible level to all the tested isolates.

### Bgt races preparation and inoculation

Local mixed races of *Bgt* were maintained on high susceptible variety o6Y86 seedlings in the greenhouse under 14 h light/10 h dark (24/18°C, 70% humidity). The single race E31, kindly provided by Dr Yining Zhou, CAAS, was maintained on high susceptible variety Sumai No.3 seedlings in the separately incubator with similar light temperature condition. Wheat seedlings at the two-leaf stage were inoculated with *Bgt*, and leaves were harvested at different time points after inoculation for gene expression analysis.

### Construction of NLR Enrichment genome DNA library of *H.villosa* and sequencing by PacBio

To accelerate *NLR* gene cloning, a new method was applied for the whole genome NLR genes development of *H.villosa,* which combined the NLR enrichment with long read PacBio sequencing offered by the RSII platform. The unique Barley NLR bait library (shared from Matthew Moscou, private communication) with 99,421 NLR baits of 100 nucleotides to capture the NLR complement from the original *Pm21*-carrying *H.villosa* accession 91C43. The construction of genomic DNA library for RenSeq and the NLR capture procedure was performed as described by Witek et al. (2016) and then sequenced on four PacBio SMRT cells.

### RNA-Seq f *H.villosa* and annotated with NLR-parser

The original *Pm21*-carrying *H.villosa* accession 91C43 was inoculated with freshly mixed local *Bgt* races and leaves were sampled at 0, 6 and 24 hours after inoculation (hai). The samples for transcriptome analysis were the mixture of equal amount of the two sample's RNA and sequenced on paired-end Illumina HiSeq 2000 platforms at BGI.

### PCR amplification for mapping of the putative *NLRs* on 6VS

All primers used for PCR are listed in Table 1, including NLR1-V and NLR2-V for physical mapping, two pairs of SNP primers NLR1-IP and NLR1-OP for co-segregation analysis. The PCR was performed in 25-μ1 reaction volumes including 1× PCR buffer, 2 mmol/l MgCl_2_, 0.15 mmol/l dNTPs, 20 ng of each primer, 100ng DNA template and 1U Taq DNA polymerase (TaKaRa, Japan). The conditions for thermal cycling involved incubation at 94°C for 3 mins, followed by 33 cycles that each involved 94°C for 45 s, 55°C for 45 s, and 72°C for 1 min. The PCR products were separated in 0.8% Agarose gels. Alignment of the amplified sequences was conducted for nucleotide variations detection using by DNAMAN 5.2.2 software.

**Table 1.**
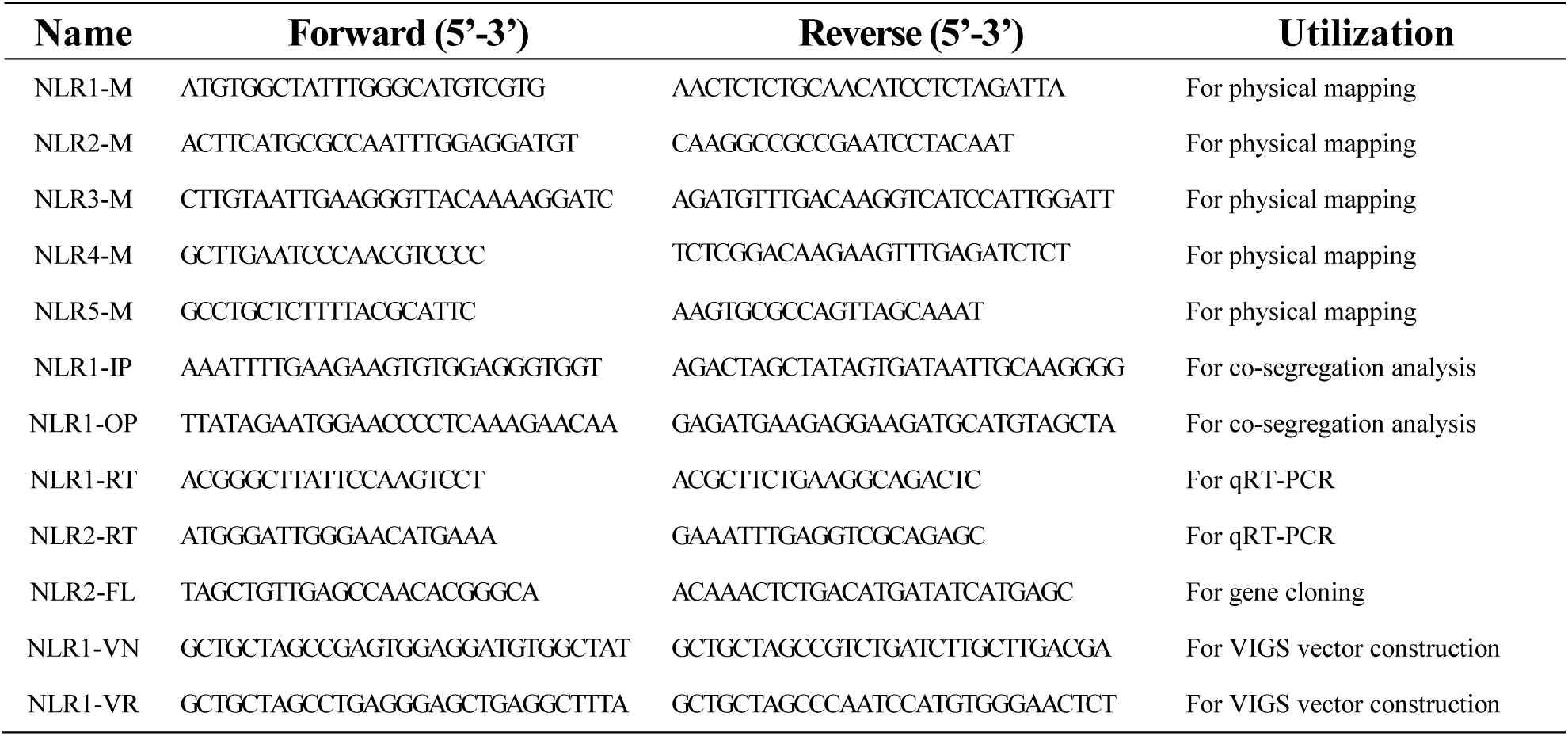
Primer sequence.

### Gene cloning and sequence analysis

Primers of NLR1-FL and NLR2-FL (Table 1) were used for cloning the full length of ORF from the cDNA of T6VS · 6AL and the six mutants. The amplified NLR1-V and NLR2-V gene fragment were ligated to pTeasy-18 vector (TaKaRa, Japan) for sequencing at Tsingke (Nanjing, China). The putative domain of the cloned genes were analyzed using the BLAST (http://www.ncbi.nlm.nih.gov/blast/), the multiple sequence alignment and the phylogenetic tree were conducted by Geneious9.1.4 software.

### Expression analysis of *NLR1-V* by qRT-PCR

Total RNA was isolated using Trizol Reagent (Invitrogen, USA) and quantified with a NanoDropTM 1000 spectrophotometer (Thermo Fisher Scientific, USA). The first-strand cDNA was synthesized using 2 μg of total RNA by the AMV reverse transcriptase (Takara, Japan) following the manufacturer's instruction. *NLR1*-RT and *NLR2*-RT (Table 1) specific primers were used for expression analysis of *NLR1-V* by qRT-PCR, and *Tubulin* gene was the internal control for normalization. The PCR reaction was performed in 25 μ1 of reaction mixture containing 1×SYB Premix Ex Taq (Takara, Japan), 0.2 μΜ of each primer, and about 30 ng cDNA per sample using the ABI Prism 7500 system (Applied Biosystems, USA). The program used was as follows: 1 min at 95°C, followed by 40 cycles at 95°C for 10 s, 60°C for 20 s, and 72°C for 40s. Three independent biological replications were performed for each treatment. The comparative 2^−ΔΔCT^ method was used for quantify relative gene expression.

### Single-Cell transient over-expression assay

The single-cell transient over-expression assay was performed as described by Shirasu et al. (1999). The over-expression vector pBI220-35s:*NLR1*-V was firstly constructed and the pAHC20 vector containing reporter GUS genes was selected for co-transformation. The two plasmids were mixed before coating of the particles with molar ratio of 1:1 in 1 μg of total DNA. The fresh seedling leaves were pre-cultured to 1% agar plates supplemented with 20mg/ml 6-BA for 1h and restoratively cultured after bombardment for 8h after at room temperature. Then, the leaves were inoculated with high-density fresh *Bgt* spores and incubated at 18°C for 48h. GUS staining cells and fungi staining with Commassie blue R250 on the leaves were observed by using Olympus BX-60 fluoroscope (Japan), subsequently. The haustorium index (percentage of GUS-staining cells with haustorium in the total GUS-staining cells interacted with *Bgt*) was calculated for each contributing at least 100 interactions, and the value represents the mean of three independent experimental repeats.

### Gene Silencing by BSMV-VIGS

Barley stripe mosaic virus induced gene silencing (BSMV-VIGS) was used to characterize the *NLR1-V* gene function in T6VS · 6AL and T6VS · 6DL. Specific primers (Table 1) were designed for amplifying the short fragment targets located on conserved NB-ARC and LRR domain coding sequencing, respectively. Then targets were reversely inserted into the γ-strain of the BMSV by replacing the *TaPDS* sequence of the BSMV:*TaPDSas* vector for construction of the recombinant silencing vectors *BSMV*:*NLR1-Vas.* The plasmid linearization, in vitro transcription and virus inoculation were performed as described by Wang et al. (2010). The second fully expanded leaves of the T6VS · 6AL and T6VS · 6DL were infected with in vitro transcribed virus *BSMV*:*NLR1-Vas,* using BSMV*:TaPDSas* and BSMV:γ as the controls, and the 5^th^ full expanded leaves with clear virus infection symptom were detached and placed on the 1% agar plates supplemented with 20mg/ml 6-BA. Then, the detached leaves were inoculated with high-density fresh *Bgt* spores and incubation at 25°C with the 14/10 h light/dark photoperiod. Photos with powdery mildew disease phenotype were taken at 7 days after *Bgt* inoculation. The bleached leaves were stained with Commassie blue for observing the fungi development in bright field by Olympus BX-60 Stereo-Fluoroscope (Japan). qRT-PCR were performed to evaluate the silencing efficiency of the target gene at transcriptional level (Primers in Table 1). At least 10 plants of each experiment had been challenged by each BSMV vector and the experiments were repeated for three times.

## Results

### 1. Screening of the full length expressed *NLRs* from *H. villosa* by RenSeq-PacBio and RNA-Seq putative located on the 6V chromosomes

The new RenSeq-PacBio approach provided us a rapid way for cloning full length disease resistance genes from the whole genome wide. From RenSeq-PacBio of *H.villosa* genome data, 107,153 3-MinFullpasses-90-MinPredicte-dAccuracy ROIs were generated with average length of 4.5 Kb and annotated with NLR-parser (NLR-specific motif alignment and search software). The ROIs containing NLRs were de novo assembled with Geneious 9.1.4, resulting in 1509 contigs that harboring 482 full length and partial NLRs after filtering the duplicated ones. The same annotation with NLR-parser was carried out using the de novo assembled RNA-Seq transcripts, and 457 transcripts harboring NLR components were found, which were deduced expressed NLR transcripts.

To explore the full length expressed NLRs of *H. villosa* putatively located on the 6V chromosome, the collinearity analysis was carried out using the public sequence database of the homeologous group 6 chromosome. Firstly, we compared the NLR complements explored by RenSeq-PacBio to the published online database IWGSC and Barley (IPK, Popseq data). Totally, 45 individual NLR-carrying contigs showing >80% identity with 6A/6B/6D/6H NLR homologous were obtained which were identified as the putative *NLRs* located on the 6V chromosome. Then, we anchored 45 full length genomic *NLRs* to the expressed *NLRs* from RNA-Seq short-read data, and 16 full length expressed *NLRs* were screened.

### 2. Identification of a new cryptic alien introgression HP33 with *Pm21* locus

T6VS · 6AL was not a suitable material to precisely locate the resistance candidate genes in the *Pm21* locus, because 6VS chromosome arm was too large and cannot recombination with 6AS chromosome arm. A new cryptic alien introgression HP33 was screened from the large irritation population of T6VS · 6AL, which showed no fluorescence signal when analyzed with GISH using the H. *villosa* genomic DNA as probe in the mitotic metaphase of root-tip cells. However, the HP33 showed resistance to powdery mildew throughout the whole growth stage, and showed polymorphic specific to *H. villosa* when analyzed with the molecular marker, it was proposed that HP33 contained the *Pm21* locus.

Compared with the resistant terminal translocation line NAU418 and resistant interstitial translocation line NAU419 in which the translocation segments can be detected by GISH (Chen et al., 2013), HP33 contained even smaller introgression segment involved 6VS. HP33 was a perfect material to precisely locate the *NLRs* genes in the *Pm21* locus.

### 3. Screening of the NLR genes in the *Pm21* locus by using HP33

Several genes have been reported as the powdery mildew resistance related gene in the 6VS, and these genes were all located on the 6VS (FL 0.45-0.58) using the resistant deletion line del.6VS-1 (FL0.58) and a susceptible deletion stock del.6VS-2 (FL0.45) (Cao et al., 2011, He et al., 2016). However, none of these genes belonged to the NLR gene and silencing of these genes could only partially compromised the resistance of T6VS · 6AL. So, after the HP33 was screened, it was tried to screen the expressed NLR genes in the introgression segment of HP33 and to find out their relationship with *Pm21*.

A number of molecular markers were developed specific to the 6VS segment introgressed in HP33, which was used for the comparative genomics analysis. Five of the expressed *NLRs* were mapped to 6VS segment by comparative analysis. Then, the primers specific to five *NLRs* genes were designed to perform PCR for physical location of them, and it was detected that only two *NLRs* could amplify single specific band in HP33 as that in 6VS chromosome (Fig.1). So, these two genes, named the as *NLR1-V* and *NLR2-V,* were exactly physical located on the *Pm21* locus. It was also found that *NLR1-V* and *NLR2-V* was two full length *NLRs* located on the same RenSeq-PacBio Hv-contig-39 (Fig.2a).

**Figure 1.**
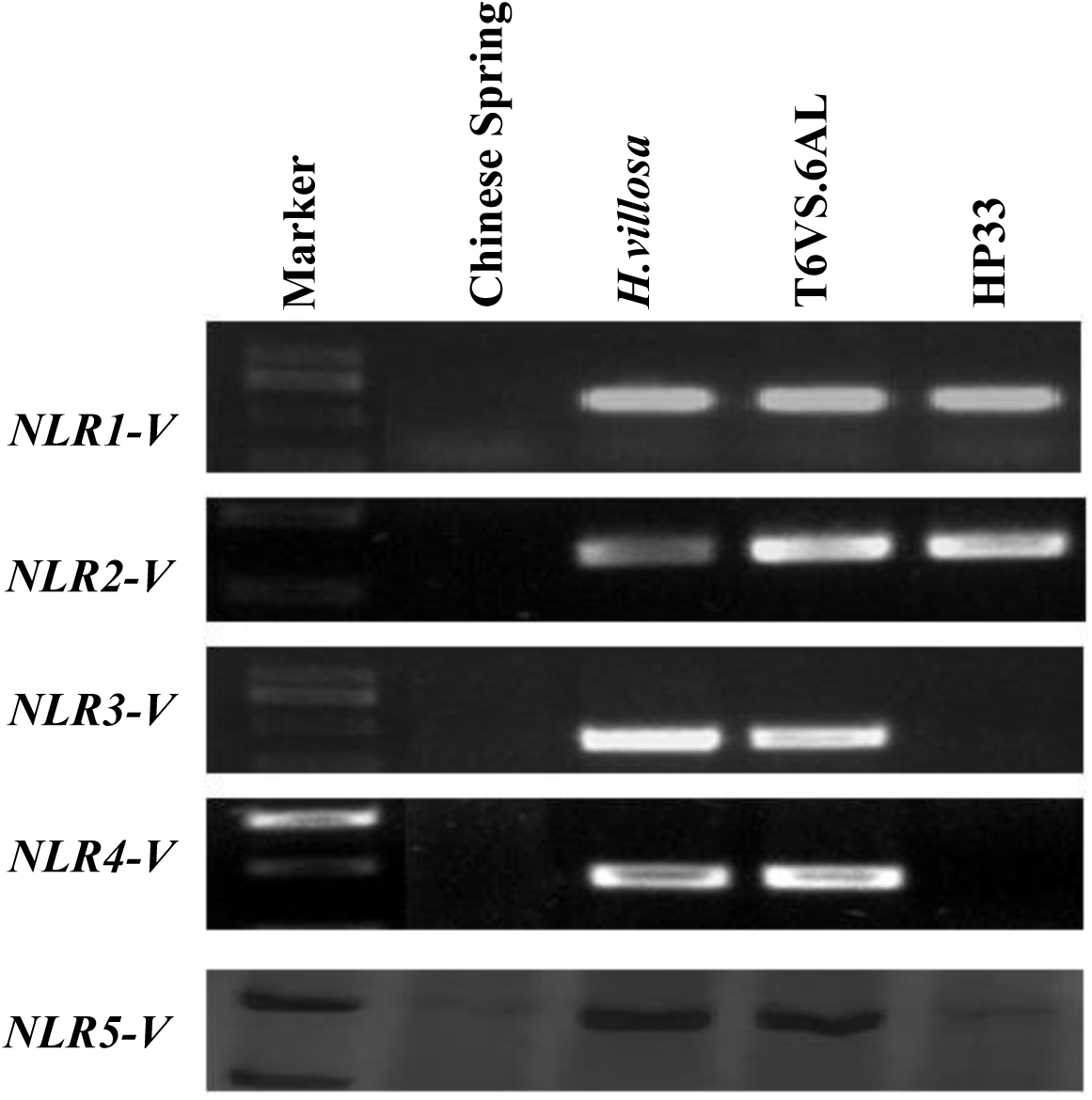
PCR-based physical location of the five expressed *NLRs.* Five expressed *NLRs,* mapped to 6VS segment by comparative analysis, were physically mapped by using four cytogenetic stocks. *NLR1-V* and *NLR2-V* were mapped to the *Pm21* locus carried in the cryptic translocation line HP33. Marker indicates the molecular marker DL2000 (TAKARA, Japan).

**Figure 2.**
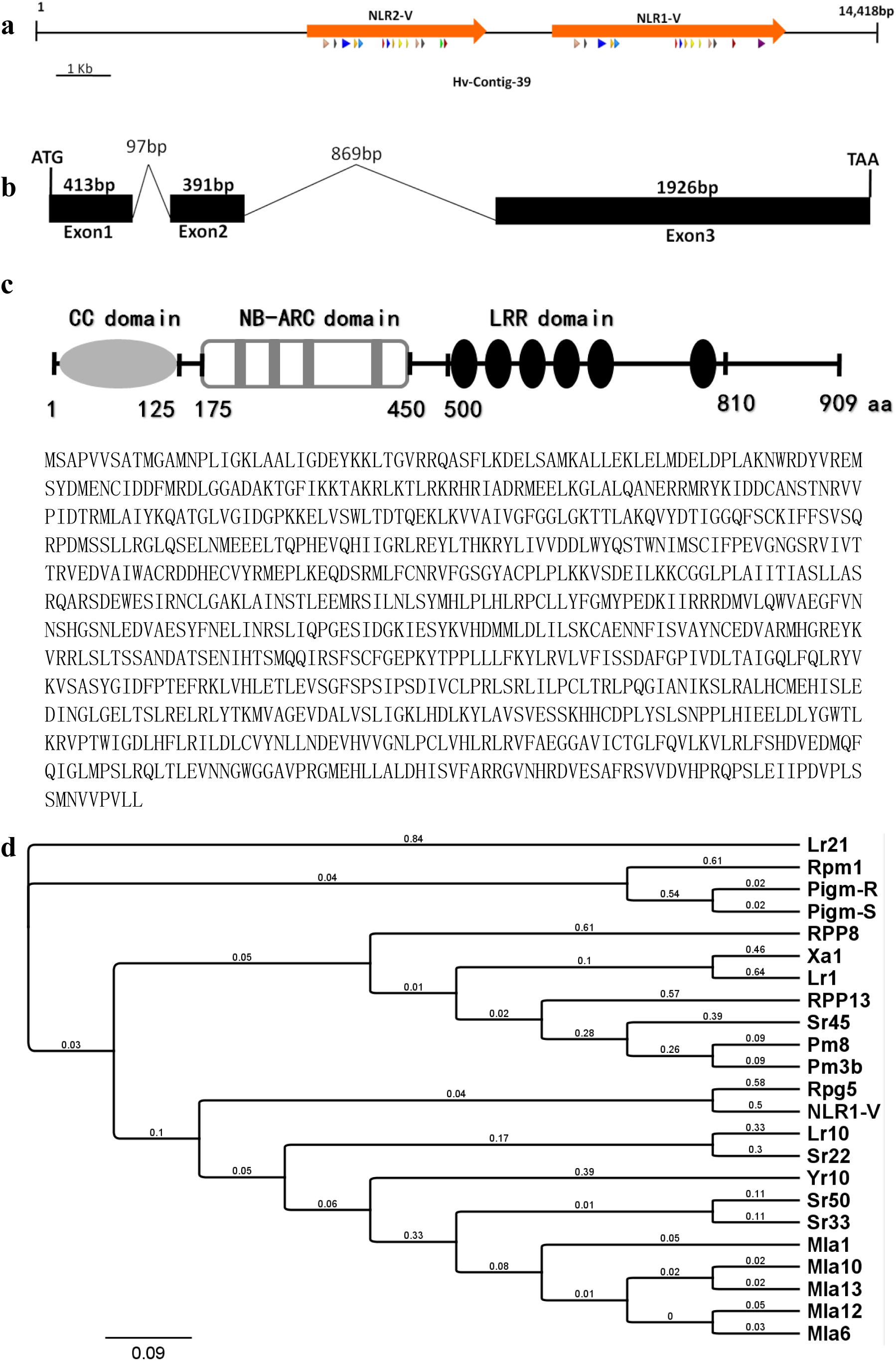
Sequence analysis of *NLR1-V* gene. a) Two paired NLR genes was predicted from RenSeq-PacBio contig39 (in length of 14,418 bp), which are named as *NLR1-V* and *NLR2-V,* respectively (showed in orange). Multiple small triangles under the predicted *NLR* genes indicate the conserved NLR motif annotated by NLR-parser software. b) Gene structure of *NLR1-V.* ORF of *NLR1-V* is determined and compared with the genomic sequence. Exons are indicated by black rectangle, and introns are indicated by lines. Sizes of them are indicated in bp. c) NLR1-V is a typical CC-NB-LRR resistance protein. The conserved domains are indicated. Sizes of them are indicated in aa. Coding protein sequence of NLR1-V was attached. d) The phylogenetic analysis of NLR1-V. Several reported R proteins were selected from hexaploid wheat, barley, rice and *Arabidopsis thaliana* for building the Neighbour-joining tree, including Mla1 [AAG37356.1, *Hordeum vulgare* subsp. *vulgare],* Mla6 [CAC29242.1, *Hordeum vulgare* subsp. *vulgare],* Mla10 [AAQ55541.1, *Hordeum vulgare],* Mla12 [AAO43441.1, *Hordeum vulgare* subsp. *vulgare],* Mla13 [AAO16000.1, *Hordeum vulgare],* Rpg5 [AGJ95079.1, *Hordeum vulgare* subsp. *spontaneum],* Sr22 [CUM44200.1, *Triticum aestivum],* Sr33 [AGQ17382.1, *Aegilops tauschii],* Sr45 [CUM44213.1, *Triticum aestivum],* Sr50 [ALO61074.1, *Secale cereale],* Yr10 [AAG42167.1, *Triticum aestivum],* Lr1 [ABS29034.1, *Triticum aestivum],* Lr10 [ADM65833.1, *Triticum dicoccoides],* Lr21 [ACO53397.1, *Triticum aestivum],* Pm3b [AAQ96158.1, *Triticum aestivum],* Pm8 [AGY30894.1, *Triticum aestivum],* Xa1 [BAA25068.1, *Oryza sativa Indica* Group], Pigm-R [APF29096.2, *Oryza sativa Indica],* Pigm-S [APF29097.1, *Oryza sativa Indica],* Rpm1 [AGC12590.1, *Arabidopsis thaliana],* RPP13 [AAF42830.1, *Arabidopsis thaliana],* RPP8 [AAC78631.1, *Arabidopsis thaliana]* and the NLR1-V.

### 4. Cloning of the *NLR1-V* and sequence analysis

Susceptible mutants were the valuable materials for the candidate gene evaluation, so after the *NLR1-V* and *NLRV-2* were physically located on the *Pm21* locus, the sequence of these two genes were preliminary compared between the T6VS · 6AL and the mutant SM-1. The specific primers (Table 1).to clone the full length of *NLR1-V* and *NLR2-V* were easily designed according to the RenSeq-PacBio Hv-contig-39 sequence. It was found that the sequences of *NLRV-2* was exactly the same in T6VS · 6AL and in the SM-1, while one SNP leading to the amino acid change was detected in the *NLR1-V* of SM-1 when compared with that in T6VS · 6AL (Sequence analysis data of *NLR2-V* gene is not shown). So, *NLR1-V* was selected as the target gene for the following analysis.

We aligned the cDNA sequences to the genomic sequences to determine the gene structure of *NLR1-V,* and it indicated that *NLR1-V* gene has three exons and two introns. The genomic sequence of *NLR1-V* spans 3,696 bp from the translation initiation (ATG) to termination codons (TAA), and the cDNA ORF sequence was 2,730 bp in length which encoded a predicted protein of 909 amino acids (Fig.2b). The encoded structure analysis showed that the NLR1-V was a typical CC-NBS-LRR resistance protein containing the conserved coiled coil, nucleotide binding site, and leucine-rich repeats domains (Fig.2c).

The phylogenetic analysis showed that Rpg5 from barley (AGJ95079.1, *Hordeum vulgare* subsp. *spontaneum*) was the most closely R protein to NLR1-V, which was reported playing critical role in barley strip rust (Arora et al., 2013). The recent reported broad-spectrum R proteins Pigm-R and Pigm-s from rice (Deng et al., 2017) involved in a distinct clade. (Fig.2d)

Expression of the *NLR1-V* gene was rapidly induced in the seedlings of resistant T6VS · 6AL during *Bgt* infection, with transcription levels peaking at 12-24h after inoculation. Similar inducible expression pattern was also observed in the *Bgt*-susceptible mutant SM-1 (Fig.3). It suggested the involvement of *NLR1-V* gene in *Pm21*-mediated *Bgt* resistance response, but the mutation in the coding region did not change the expression of the *NLR1-V* gene.

**Figure 3.**
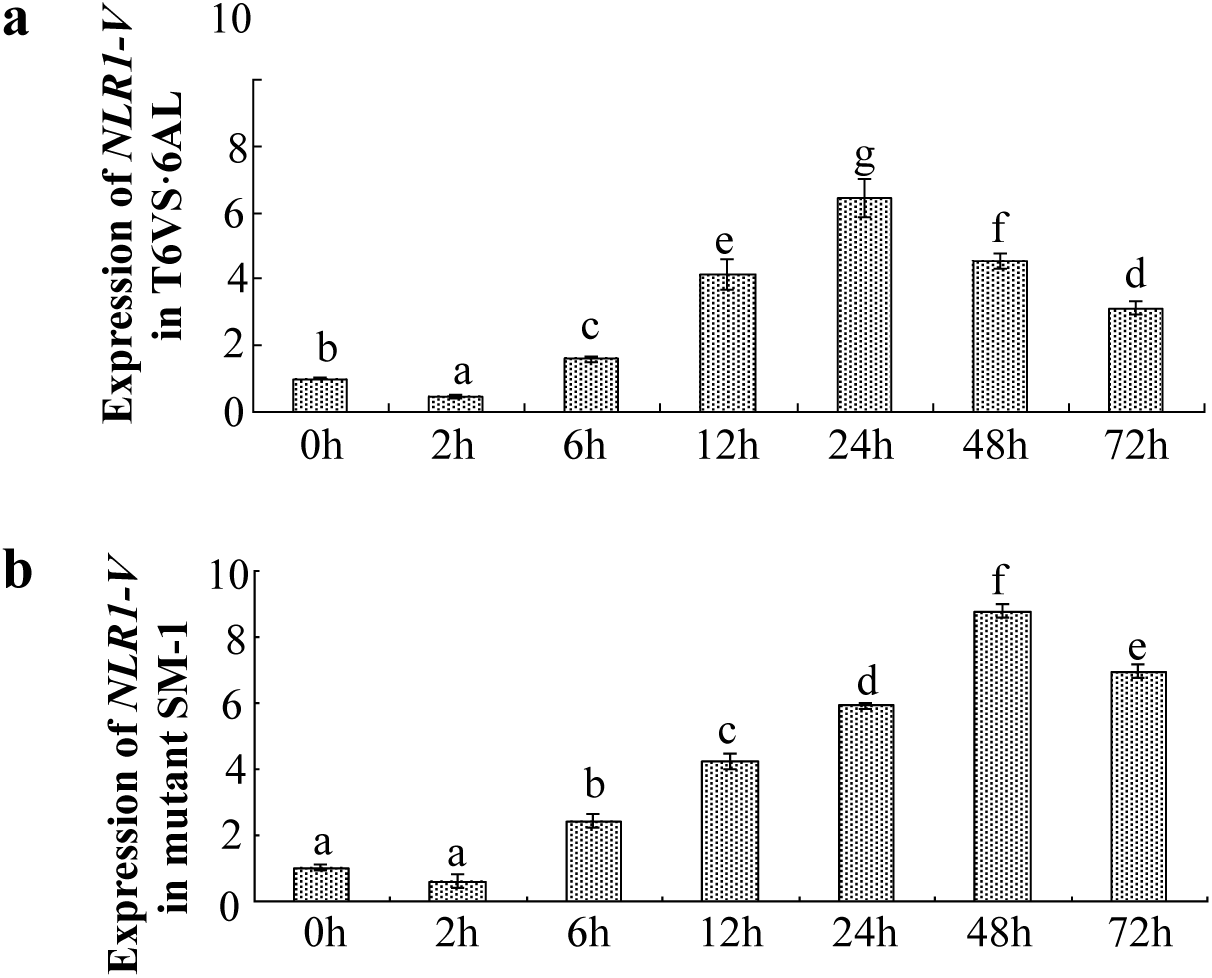
Expression pattern of *NLR1-V* in the leaves of T6VS · 6AL and the mutant SM-1 before and after *Bgt* inoculation. Expression level of *NLR1-V* was up-regulated at 6 hours after *Bgt* inoculation and then decreased at the later inoculation stage both in T6VS · 6AL (a) and in the SM-1 (b). The mutation of *NLR1-V* did not change its expression pattern.

### 5. Comparison of *NLR1-V* between the wild T6VS · 6AL and its six independent mutants

In the previous study, six of lose-of-function independent mutants of T6VS · 6AL were identified by EMS-treatment, which showed *Bgt*-susceptible symptom when challenged with the local mixed *Bgt* races and single race E31. Meanwhile, the wild-type T6VS · 6AL showed the high resistant to *Bgt* (Fig.4a and 4b). The SNP in the coding region has been detected in the SM-1, so the *NLR1-V* was also cloned from the remaining five mutants to compare the difference of the sequences. It was exciting to find that there was SNP between the *NLR1-V* of T6VS · 6AL and that in the all mutants. The SNP in SM-1, SM-2, SM-3 occurred in the NB-ARC region, and the nucleotide change from C to T leading to the amino change from P to S. In SM-4, SNP also occurred in the NB-ARC region, and the nucleotide changed from G to A leading to the amino change from A to T. In SM-5, SNP also occurred in the LRR region, and the nucleotide changed from C to T leading to the amino change from S to L. In SM-6, SNP occurred in the NB-ARC region, and the nucleotide changed from G to A leading to the leading to the early termination and missing of the whole LRR region (Fig.4c.

SM-1 was the first identified mutant in 2007, and the F2 population was developed by cross the SM-1 with its wild type T6VS · 6AL. After the SNP was detected in the SM-1, the primers were designed according the SNP site to determine whether the SNP was linked with the mutant phenotype (Primers in Table 1). It was found that all the homozygous resistant plants showed the same polymorphic as in T6VS · 6AL, and all the homozygous susceptible plants showed the same polymorphic as in the mutant SM-1(Fig.4d). So, the SNP in the *NLR1-V* was responsible for the resistance lose.

**Figure 4.**
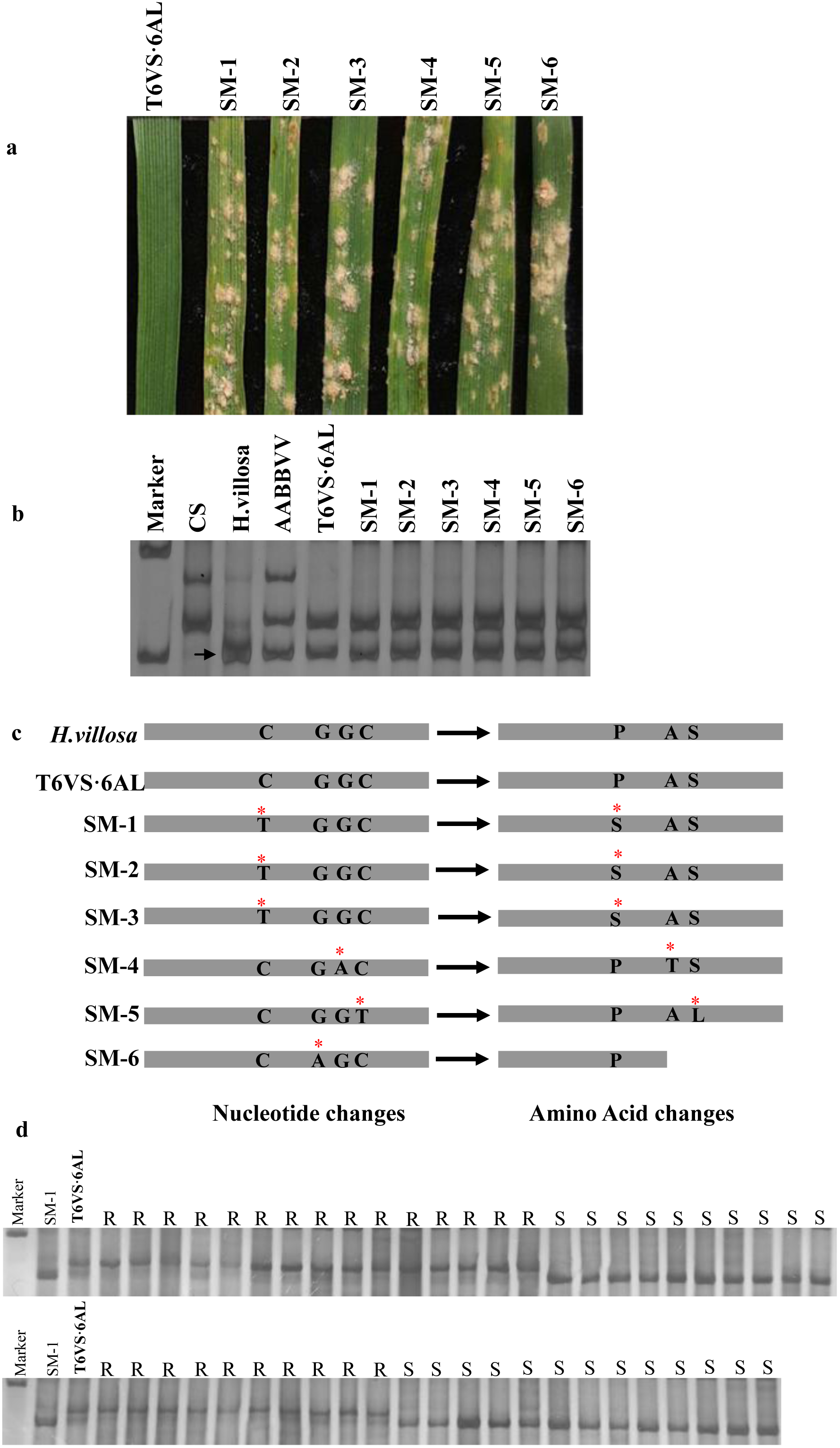
*NLR1-V* comparison between the susceptible mutants and its wild-type T6VS · 6AL. a) Six powdery mildew susceptible mutants, SM-1 to SM-6, screened from the EMS treated T6VS · 6AL population. b) Detection of genetic background of the mutants and the wild-type T6VS · 6AL, and the arrow indicates the polymorphism specific to 6VS chromosome arm. c) Changed nucleotide acids and the corresponding amino acids in the *H.villosa,* wild T6VS · 6AL and six susceptible mutants. Red * indicates the mutated point. d) Linkage analysis of the SNP molecular marker, developed according to the mutated point in the SM-1, with the phenotype of the individual plants in the F_2_ population crossed by T6VS · 6AL and SM-1.

### 6. Functional analysis of *NLR1-V* with transient assay

*NLR1-V* gene was over-expressed in wheat leaf using the transient single-cell and *Bgt* interaction assay system (Shirasu et al., 1999) to detect its potential disease resistance function. As the control transient expressed GUS gene in SM-1, the Haustorium Index (HI) was as high as 66.67%, while when the *NLR1-V* was co-transformed with GUS, HI was decreased to 34.58% which showed no significant difference from the HI value of 36.64% in T6VS · 6AL over-expressing GUS only. But, over-expression of the mutated *NLR1-V* gene from SM-1 in the leaf epidermal cells of SM-1, the HI value was 68.44% with no significant change compared with the HI value of SM-1 transformed with GUS control 66.67% (Fig.5). These results indicate that the *NLR1-V* gene could recover the resistance of SM-1, and positively contribute to the powdery mildew resistance by preventing the haustorium formation of *Bgt.*

**Figure 5.**
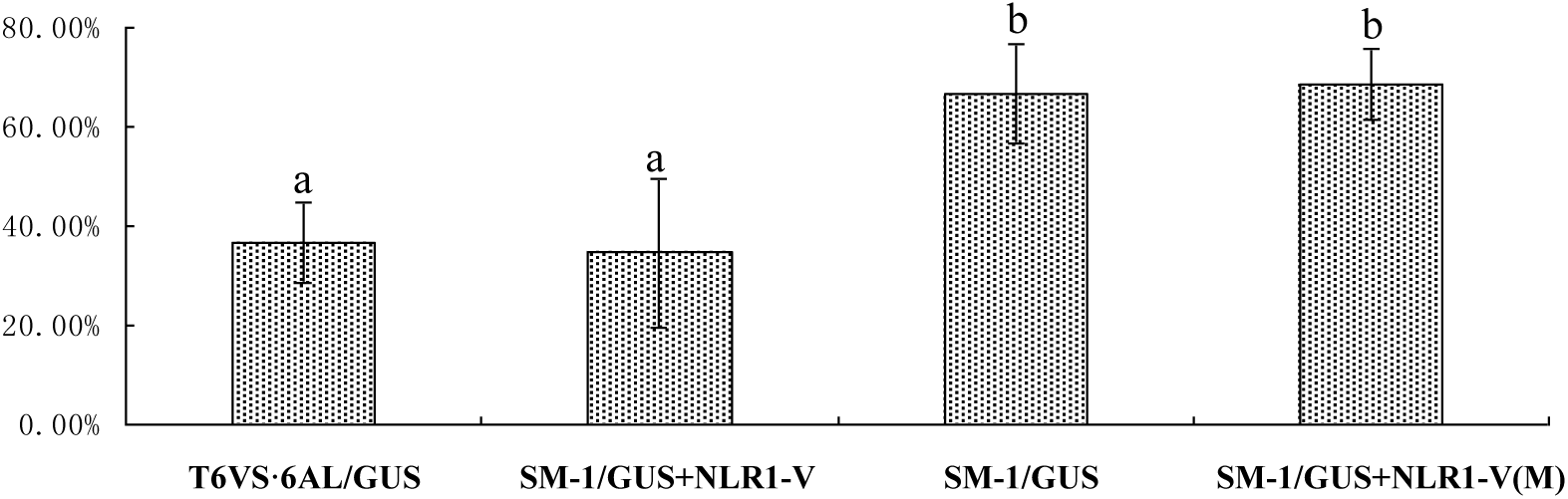
Quantitative assay of Haustorium Index (HI) in SM-1 and T6VS · 6AL. HI was determined in the susceptible mutant SM-1 transient over-expression of *NLR1-V*+GUS or over-expression of the mutated gene *NLR1-V(M)+GUS,* using the HI in the SM-1 and T6VS · 6AL transient over-expression of GUS as the controls. *NLR1-V* gene, but not the mutated gene *NLR1-V(M),* could recover the resistance of SM-1. a, b indicate significantly differences using the one-way ANOVA LSD analysis (P<0.05).

### 7. Silencing of the *NLV1-V* by BSMV-VIGS in T6VS · 6AL and T6VS · 6DL

We performed Barley stripe mosaic virus induced gene silencing (BSMV-VIGS) to characterize the function of *NLV1-V* gene in the *Bgt*-resistant T6VS · 6AL, to provide evidence supporting that *NLV1-V* gene is essential for the *Bgt* resistance of the T6VS · 6AL. For the *NLV1-V* gene VIGS in the T6VS · 6AL, both of NB-ARC domain-specific and LRR domain-specific sequences were designed as the silencing target. Both the two group of the *NLV1-V* gene silencing individuals showed susceptible to *Bgt*, as we could clearly observe generation of a large number of powdery mildew colony on the leaves. However, no spore could be observed on the leaves of Mock, empty vector BSMV:γ and *BSMV*:*PDSas* control (Fig.6a1 and 6b1). The observation of fungal development stained with Commassie blue under the microscope also showed the full development of the fungal on the *NLV1-V* gene silencing leaves (Fig.6a3 and 6b3), but limited development of the fungal on the controls (Fig.6a2 and 6b2). The expression of innate *NLV1-V* gene was check by qRT-PCR, which revealed the VISG system worked high efficiently with 90-98% decreasing (Fig.6c). The results indicated that silencing of *NLR1-V* could compromise the resistance of T6VS · 6AL completely. The BSMV-VIGS was also performed in the leaves of T6VS · 6DL with powdery mildew resistance gene *PmV* on the 6VS chromosome arm, and it found that silencing of *NLR1-V* could also decrease the resistance of T6VS · 6DL dramatically (Fig.6d1 and 6d2).

**Figure 6.**
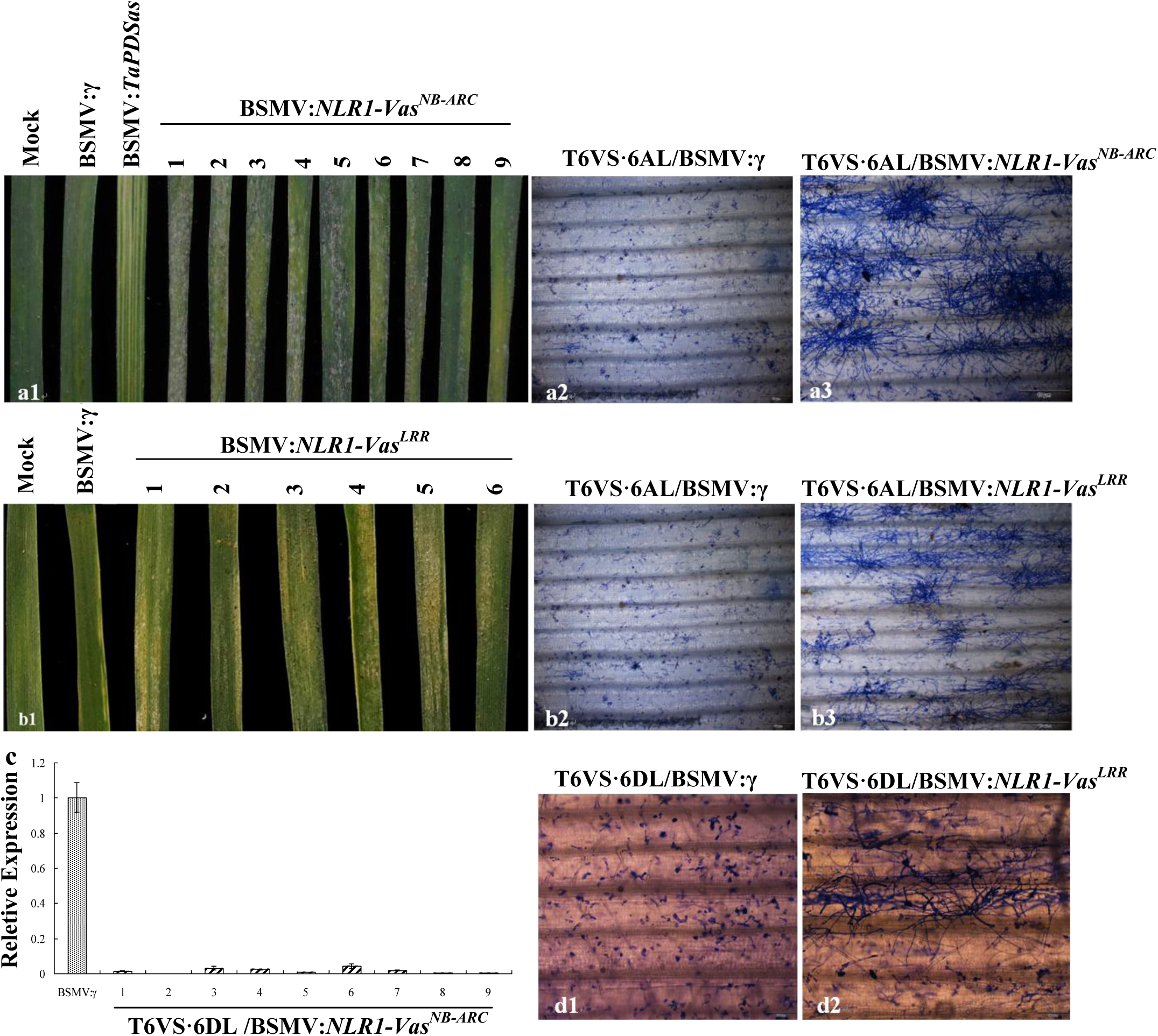
Disease resistance function of *NLR1-V* analysis using BSMV-VIGS. a) When the *NLR1-V* was silenced by VIGS targeting to NB-ARC region in T6VS · 6AL, the resistance was compromised completely. a1) Representative photographs were taken 7 days after *Bgt* inoculation, and the *Bgt* colony could be observed on the BSMV:*NLR1-V*as^NB-ARC^ infected leaves while not the Mock, empty vector BSMV:γ and BSMV:*TaPDS*as infected leaves. a2-a3) Under the microscope, *Bgt* could not developed in the BSMV:γ infected leaves of T6VS · 6AL (a2), but could developed normally in the BSMV:*NLR1-V*as^NB-ARC^ infected leaves (a3). b) When the *NLR1-V* was silenced by VIGS targeting to LRR region in T6VS · 6AL, the resistance was compromised completely. b1) Representative photographs were taken 7 days after *Bgt* inoculation, and the *Bgt* colony could be observed on the BSMV:*NLR1-V*as^LRR^ infected leaves while not the Mock, empty vector BSMV:γ and BSMV:PDSas infected leaves. b2-b3) Under the microscope, *Bgt* could not developed in the BSMV:γ infected leaves of T6VS · 6AL (b2), but could developed normally in the BSMV:*NLR1-V*as^LRR^ infected leaves (b3). c) qRT-PCR analysis of *NLR1-V* expression level in both BSMV:γ and BSMV:*NLR1-V*as^NB-ARC^ challenged individuals. d) When the *NLR1-V* was silenced by VIGS targeting to LRR region in T6VS · 6DL, the resistance was decreased dramatically. Histology observation of the fungi development on the leaves from T6VS · 6DL individuals challenged with BSMV:γ (d1) and BSMV:*NLR1-V*as^LRR^ (d2), respectively.

## Discussion

### The approach of mutagenesis, RenSeq, PacBio and cytogenetic stocks development were combined in the exploration of the *NLR1-V* gene

The wild species usually confer high resistance to the serious diseases and tolerance to the frequent abiotic stresses, so they provide rich gene resources for the improvement of the cultivated crop. In wheat, the elite genes from several wild species have been successfully used in crop production. However, due to the evolutionary distance between the wild species and common wheat, chromosome containing the elite gene cannot easily recombination with the chromosome of wheat, which leading to the difficulty of the wild gene cloning. So, development of the cytogenetic stocks containing the target gene but with less redundancy genes seems to be very important to precisely locate the candidate genes. In this study, a new cryptic alien introgression HP33, which contains a very small 6VS segment which can only be detect by molecular markers play a very key role to fine mapping of the candidate NLR genes in the *Pm21* locus.

In recent years, with the development of the sequencing technology, R gene cloning in wheat was pushed forward quickly by combine mutagenesis and RenSeq, for example, Sr22 and Sr45 resistant to stem rust in wheat were clone using this approach (Steuernagel et al., 2016). With the emerging of the new sequencing technology PacBio which can get longer assembled sequences, the R genes cloning were tried by combing RenSeq and PacBio, for example, Rpi-amr3 resistance to potato late blight was clone using this approach (Witek et al., 2016). The lack of reference genomic sequence makes the full length gene screening particularly difficult in *H. villosa,* so the RenSeq-PacBio is a perfect way to construct the database of the full-length NLR genes. In addition, a number of susceptible mutants of provide valuable materials to validation of the target genes, which is extremely feasible when the genetic map could not be constructed due to the non-recombination. So, the *NLR1-V* cloning approach was combined by mutagenesis, RenSeq, and PacBio, and this approach provide valuable examples for R gene cloning from other wild species.

### The *NLR1-V* was a potential candidate gene of *Pm21*

Several genes were found to be related to the resistance mediated by *Pm21*, however, silencing of these five genes can not compromised the resistance of T6VS · 6AL completely (Cao et al., 2011; He et al., 2016). *Stpk-V* was previously cloned by our group, and in the following research it was found that the transgenic plants over-expression of *Stpk-V* can resistant to 23 testes *Bgt* isolates but susceptible to *Bgt* isolate E31, however, the T6VS · 6AL could resistant to all of the 24 tested isolates. So, it was proposed that there should be another gene which perhaps plays resistance role along with *Stpk-V* or regulated *Stpk-V* directly. *NLR1-V* was found to be mutated in all the screened susceptible mutants, and it can recover the complete resistance in the mutant. As a CC-NBS-LRR gene, the overwhelming majority class of cloned R genes in cereals, *NLR1-V* was a potential candidate gene of *Pm21.*

### The NB-ARC and LRR domain were both critical for the function of *NLR1-V*

It was reported that the LRR domain usually recognize the effectors of pathogen (Dangl and Jones 2001), which can make the structure changes of the NB-ARC domain leading to activation of the signal transduction (Inoue et al., 2013). However, in some cases, the NB-ARC domain could also recognize the pathogen effector, such as N gene of tobacco could interact with pathogen effector molecule p50 (Burch-Smith et al., 2007). In this study, the mutation of the LRR domain or the NB-ARC domain could compromised the resistance of the T6VS · 6AL completely, which implied that the both domains were critical for the function of *NLR1-V.*

NLR genes usually induced cell death in the resistance pathway, in some cases the resistance was dependent on the cell death induced by R genes, however, in other cases the resistance was independent on the cell death (Chang et al., 2013). It was also reported that the domain inducing cell death was sometimes different from the domain inducing resistance. Cell death could be observed obviously in the leaf of T6VS · 6AL 24 hours after *Bgt* inoculation, so it was interesting to study whether cell death disappeared when the resistance lost in the susceptible mutants. It will be helpful to evaluation the relationship between the cell death and the disease resistance mediated by *NLR1-V.*

## Acknowledgements

We highly appreciate Dr. Kamil Witek (The Sainsbury Laboratory, UK) for kind help in accomplishing RenSeq-Pacbio library construction and database building, Dr. Matthew Moscou (The Sainsbury Laboratory, UK) for kind help in sharing the barley baits library and analysis of the transcriptom data. We thank Prof. Jonathan D. G. Jones (The Sainsbury Laboratory, UK) and Brande Wulff (John Innes Centre, UK) for kind suggestion of this work. This work was supported by Natural Science Foundation of China (Grant No. 31671685, 31471489), the Important National Science & Technology Specific Projects of Transgenic Research (Grant No. 2014ZX0800202B-002), the Fundamental Research Funds for the National Central Universities (Grant No. KYZ201601, KYYJ201602, KYZ201401).

